# Neonatal Enteric Infection Disrupts the Microbiota-Gut-Brain Axis Through Pattern Recognition Receptors and Altered Neuroimmune Signaling

**DOI:** 10.64898/2026.01.14.699546

**Authors:** Jungjae Park, Olivia Orahood, Maithili Banginwar, Amy Andrade, Gatha Pore, Kristina Sanchez, Michael Cremin, Jelissa Reynoso-Garcia, Sarah Lee, Ingrid Brust-Mascher, Mariana Barboza, Carlito B. Lebrilla, Taeseon Woo, Kara G Margolis, Colin Reardon, Mélanie G Gareau

## Abstract

Early-life enteric infection can have long-lasting effects on the microbiota–gut–brain (MGB) axis. Using a neonatal Enteropathogenic *Escherichia coli* (EPEC) model, we show that intestinal epithelial cell (IEC) NOD1 signaling coordinates mucosal immunity, barrier repair, and neuroimmune outcomes throughout early development and into adulthood. Neonates infected at postnatal day (P) 7 exhibited ileal inflammation, as demonstrated by increased expression of inflammatory cytokines (*Il1β*, *Il6*, *Il12*, *Il22*), chemokines/chemokine receptors (*Ccl2*, *Cxcl1*, *Ccr2*), and barrier-repair genes (*Muc2*, *Slc26a3*), with increased monocyte/macrophage infiltration and reduced epithelial proliferation in WT mice that was blunted in Nod1^ΔIEC^ mice. Neonatal infection of WT mice induced persistent defects into adulthood (P56), including increased intestinal permeability, sustained inflammatory/repair signatures, hippocampal inflammation, altered neurogenesis, and impaired recognition memory, which were largely absent in Nod1^ΔIEC^ mice, establishing a crucial role for IEC NOD1 as a determinant of long-term MGB remodeling. Microbially derived ligands of NOD2, muropeptides, isolated from probiotic *Lactobacillus* species attenuated EPEC-induced mucosal inflammation and chemokine induction without altering bacterial burden, demonstrating NOD2 host-directed immunomodulation. Together, these findings identify an important role for NOD-dependent signaling axis in the gastrointestinal tract that links early-life infection to enduring gut-brain dysfunction and reveals probiotic-derived muropeptides as candidate microbial therapeutics.

## INTRODUCTION

Early-life enteric infections are a major global health challenge, contributing significantly to childhood morbidity and mortality and exerting developmental consequences that extend well beyond acute diarrheal illness.^1^ Enteropathogenic *Escherichia coli* (EPEC), a non-invasive, attaching-and-effacing bacterial pathogen, remains a leading cause of pediatric diarrheal disease in children under five years of age worldwide.^2^ EPEC delivers effector proteins into intestinal epithelial cells (IEC) using a type 3 secretion system (T3SS), which mediates inflammatory signaling cascades and subsequent disruptions to mucosal barrier function.^3,4^ Importantly, these infections often coincide with critical time periods in postnatal development, when the gut microbiome, immune system, and nervous systems are rapidly maturing. Stressors, including bacterial infection, during these developmental stages may therefore make the host particularly vulnerable to long-term physiological and neurodevelopmental perturbations.^5,6^

Mounting evidence suggests that neonatal enteric infections can disrupt the developing microbiota-gut-brain (MGB) axis, a bidirectional communication network linking the gastrointestinal (GI) tract to the central nervous system (CNS) via neural, immune, and endocrine pathways.^7,8^ Perturbations to any of these routes during early life can have persistent effects on intestinal barrier integrity, immune homeostasis, and neurobehavioral trajectories. We previously reported that neonatal EPEC infection disrupts colonization of the gut microbiome and promotes systemic and neuroinflammation.^8^ While the acute effects of EPEC infection, such as dehydration, malnutrition, and stunted growth, are well-characterized^9^, the mechanisms linking neonatal enteric infection to persistent gut-brain dysfunction remain poorly defined. Acute GI inflammation and barrier disruption caused by neonatal infection^8,10–13^, may interfere with the maturation of the enteric nervous system (ENS)^14^ and neurodevelopment within the CNS, which may reprogram host physiology throughout life.

Innate immune signaling is a key mediator of host-microbe interactions during early life, particularly at mucosal surfaces, where pattern recognition receptors (PRRs) shape host-microbial immune responses.^15^ Among PRRs, the Nod-like receptor (NLR) family of intracellular PRRs are vital to innate immune activation. Nucleotide-binding oligomerization domain 1 (NOD1) plays a critical role in sensing the microbial-derived cell wall component peptidoglycan (PGN) meso-diaminopimelic acid (meso-DAP)-containing motifs found in Gram-negative organisms, including EPEC.^16,17^ Highly expressed in IECs, NOD1 activation triggers the nuclear factor-κB (NF κB) or mitogen-activated protein kinase (MAPK) pathways to induce pro-inflammatory or reparative responses in the GI tract following infection.^16–18^ Intestinal NOD1 activity further appears to shape neuroimmune responses ^19–22^, and stress-associated behavioral regulation^19^, suggesting a potential role in the MGB axis signaling.

Beyond mucosal immunity, PGN sensing pathways have also been implicated in neural development and behavior. Mice lacking PGN-sensing receptor protein 2 (Pglyrp2) exhibited sex-specific differences in social behavior and brain-derived neurotropic factor (*Bdnf*) expression, which is critical to regulation of brain function.^22^ We previously showed that Nod1/Nod2 double knockout (NodDKO) mice exhibit stress-induced behavioral impairments following exposure to acute water avoidance stress in adulthood.^19^ This effect was specifically localized to IEC NOD1 expression, identifying the gut epithelium as a potential interface between microbial recognition and neurobehavioral outcomes. Here, we investigated whether IEC NOD1 governs long-term MGB axis disruption following neonatal EPEC infection. Using IEC-specific NOD1 knockout (Nod1^ΔIEC^) mice, we found that neonatal EPEC exposure caused ileal inflammation, immune cell recruitment, intestinal permeability deficits, coupled with hippocampal inflammation, impaired neurogenesis, and cognitive dysfunction that were dependent on IEC NOD1. Finally, we identified the potential of probiotic-derived muropeptides—bioactive PGN fragments—as a microbial intervention to mitigate infection-induced pathology through host-directed immunomodulation. Our studies reveal a critical role for NOD1/2 in mediating the gut–brain communication following neonatal enteric infection and identify probiotic muropeptides as a potential therapeutic strategy.

## MATERIALS AND METHODS

### Mice

All procedures were approved by the Institutional Animal Care and Use Committee (IACUC #23455) at the University of California, Davis. IEC (Vil.Cre^+^) Nod1^f/f^ conditional knockout mice (Nod1^ΔIEC^) and wild type (WT) littermate controls (Vil.Cre^-^) on a C57BL/6 background were bred in house.^19^ Mice were weaned at P21 and housed by sex in cages under a 12-hour light/dark cycle with *ad libitum* access to water and food. Mice were euthanized by CO_2_ asphyxiation and cervical dislocation. A subset of mice was anesthetized using isoflurane, perfused with phosphate-buffered saline (PBS), and fixed with 4% paraformaldehyde for confocal imaging studies.

### EPEC

Bacterial cultures (O127:H6 strain E2348/69) were prepared from glycerol stocks maintained at -80°C. Cultures were grown in 10 ml of Luria Bertani (LB) broth at 37°C in a shaking incubator overnight until logarithmic growth was achieved, as determined by OD600 ∼ 0.6.

### Experimental Design

#### EPEC infection

Mice (male/female) were infected with 50 μL of EPEC (10^5^ CFU, serotype O127:H6 strain E2348/69) or vehicle control (LB) at P7 by orogastric gavage. In a subset of mice, serum, feces, whole brain, and distal ileum were collected at 7 days post-infection (DPI) for qPCR, histology and CFU assessments. EPEC colonization and clearance were monitored by CFU counts starting at P14 until clearance of the pathogen at P28.

At P14, acute ileal inflammation and infection-induced GI pathology were assessed via qPCR and immunofluorescence imaging, flow cytometry, and histology. In early adulthood, (P42-56 [6-8 weeks]), behavior was assessed via a battery of three validated tests, the novel object recognition (NOR) task, the light/dark (L/D) box, and the open field test (OFT).

Inflammation (ileum and hippocampus) was assessed via qPCR, hippocampal neurogenesis was assessed using qPCR and immunofluorescence imaging, and corticosterone levels measured via ELISA. (**Fig 1A**)

**Figure 1.**
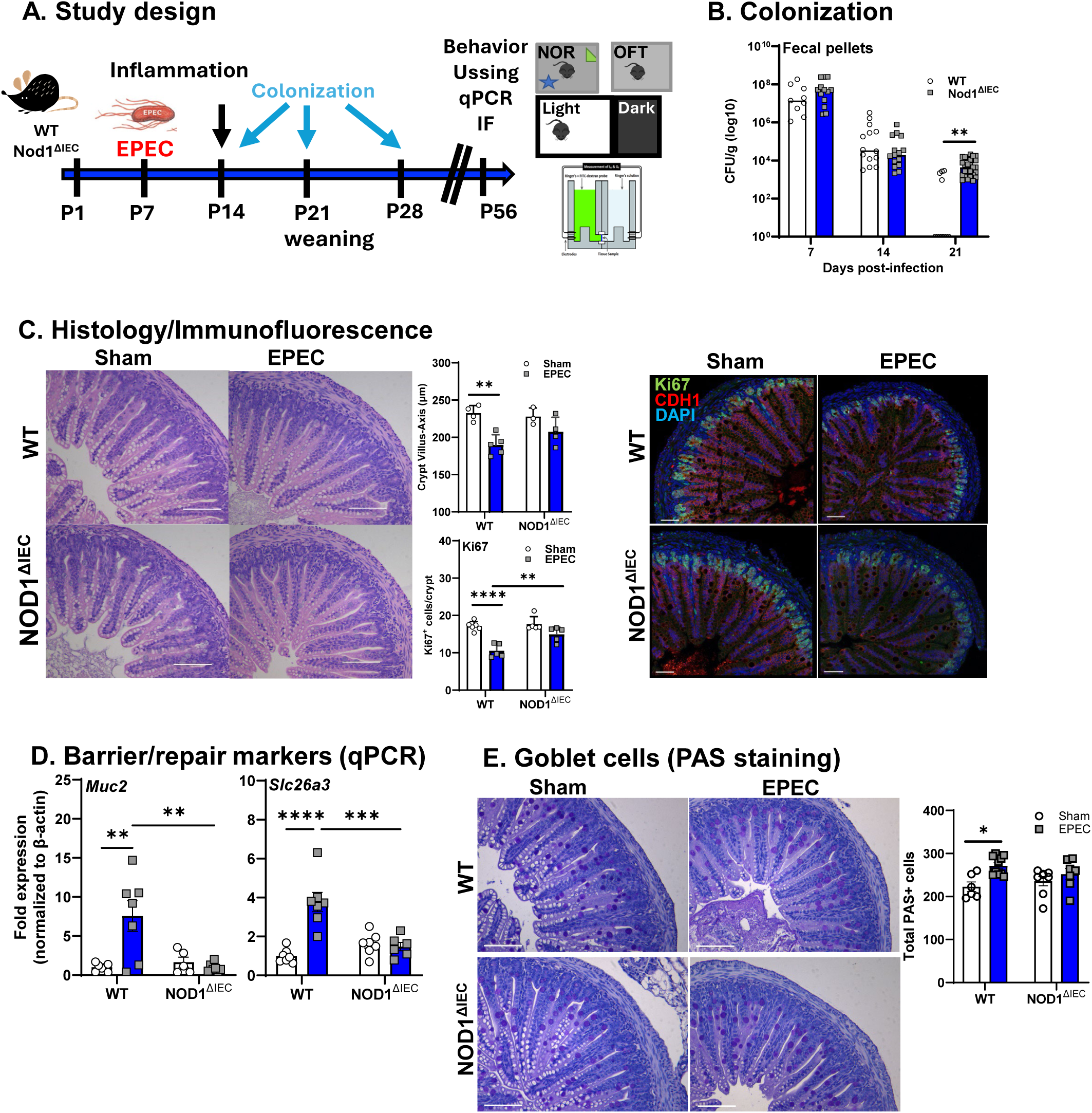
Intestinal epithelial NOD1 is required for efficient clearance of neonatal EPEC infection and drives altered intestinal epithelial cell kinetics. Study design of neonatal EPEC infection in wild type (WT) and NOD1 knockout (Nod1^ΔIEC^) mice (**A**). Bacterial colonization levels (CFU/g feces, log10 scale) were assessed at 7-, 14-, and 21-days post-infection in WT and Nod1^ΔIEC^ neonatal mice following oral EPEC infection (**B**). Epithelial cell proliferation was assessed by H&E, performed using cross sections of the distal ileum from WT and Nod1^ΔIEC^ neonatal mice following sham or EPEC infection were stained with crypt-villus axis measured, coupled with immunofluorescence performed using Ki67 (green), CDH1 (red) and DAPI (blue) to quantify proliferative cells at P14 (**C**). qPCR was used to measure epithelial barrier and repair markers (Muc2 and Slc26a3) (**D**) and PAS staining was used to quantify goblet cells (**E**). N=8-20 (colonization), N=4-6 (histology/IF), N=6-8 (qPCR); mean ± SEM; *p<0.05, **p < 0.01, ****p<0.001 by 2-way ANOVA with Bonferroni correction. (Scale bar = 50 μm)

### qPCR

Tissues were collected and stored in TRIzol (Invitrogen) at -80°C until RNA extraction following the manufacturer’s protocol. Following extraction, RNA concentration and quality were measured using a NanoDrop spectrophotometer, and then cDNA synthesis was performed using the iScript cDNA synthesis kit (BioRad, Hercules, CA). Analysis of mRNA expression was determined by qPCR using PowerSYBR master mix (Applied Biosystems) on QuantStudio 6 Flex qPCR machine (Applied Biosystems, Waltham, MA) based on β-actin (housekeeping gene) expression using the 2^-ΔΔCT^ method.^23^ Data are presented as fold change relative to β-actin and normalized to WT sham mice. (**Table 1**. Primers for qPCR)

### Ussing chambers

Studies were performed as described previously.^8,24^ Briefly, segments of distal ileum were collected and immediately placed in cold oxygenated Ringer’s buffer (pH 7.35 ± 0.02) containing (in mM): 115 NaCl, 25 NaHCO₃, 1.2 MgCl₂, 1.25 CaCl₂, and 2.0 KH₂PO₄. Intestinal sections were cut open along the mesenteric border, gently rinsed to remove luminal contents, and mounted in Ussing chambers (Physiologic Instruments, San Diego, CA), exposing 0.1 cm² of epithelial surface area to 4 ml of the Ringer’s buffer at 37 °C. Glucose (10 mM) was added to the serosal chamber for metabolic support, while the mucosal chamber received an equimolar concentration of mannitol (10 mM) to maintain osmotic balance. Agar-salt bridge electrodes connected to a voltage clamp system were used to measure transepithelial short-circuit current (Isc [μA/cm^2^]), an indicator of active ion transport, and potential difference after a 15-minute equilibration period, using Acquire & Analyze II software (Physiologic Instruments). Transepithelial conductance (G [mS/cm²]), was calculated as an indicator of tight junction ionic permeability. Macromolecular permeability was assessed by adding 400 µg/mL FITC–dextran (4 kDa, Sigma-Aldrich) to the mucosal chamber and serosal samples (200 µl) taken every 30 minutes for 2 hours, and FITC concentrations were quantified using a fluorescence plate reader (Synergy H1, BioTek Instruments, Inc).

### Histology

#### Hematoxylin and Eosin (H&E)

Neonatal (P14) and adult (P56) ileum was collected and fixed in 10% formalin. Tissue was transferred to 70% IPA for dehydration after a minimum 48h post-fixation followed by processing and embedding in paraffin. Tissue sections, 5 µm thick, were transferred to positively charged slide and H&E staining was performed. Slides were assessed by light microscope (Nikon Eclipse E600) to determine histopathology and villus-crypt length.

#### Periodic Acid Schiff (PAS)

PAS staining was used to identify and enumerate the total number of goblet cells in neonatal ileal tissue sections. Briefly, slides were deparaffinized (Histoclear), rehydrated, and incubated in Periodic Acid (5 minutes in 1 g/dL). Slides were subsequently incubated in Schiff’s reagent (15 minutes), followed by washing in running tap water and counterstaining with hematoxylin (1 minute). Light microscopy was used to determine the total number of goblet cells, by manual counting in FIJI (ImageJ) in 2-3 representative sections per mouse and averaged.^25^

### Immunofluorescence

#### Epithelial cell proliferation

To determine the impact of infection on IEC proliferation at P14, ileal tissue (formalin-fixed, paraffin-embedded) was immunostained using a mouse-on-mouse immunodetection kit (Vector Labs).^8^ Proliferative cells in the crypt were stained using the primary antibody rabbit anti-Ki67 [LS-C141898; Lifespan Biosciences], and secondary antibody Alexa Fluor 555 goat anti-rabbit [A21428; Life Technologies] and co-stained with E-Cadherin (CDH1) to identify IEC, using the mouse anti-CDH1 marker [CM1681; ECM Biosciences] and streptavidin-Alexa Fluor 647 [S32357; Invitrogen] marker. 4′,6-diamidino-2-phenylindole (DAPI) was used to stain nuclei (**Table 2**). Slides were imaged using confocal microscopy with a 40x objective (SP8; Leica) and a 0.7 µm step size. Following acquisition, images were deconvolved (Huygens), with overlapping tiles stitched, and analyzed using Imaris software to automatically quantify Ki67^+^CDH1^+^ cells/crypt and measure the crypt-villus axis.

#### Hippocampal neurogenesis

Mice were anesthetized (isofluorane) and then perfused with 4% paraformaldehyde (PFA). Following perfusion, brains were collected and stored in 4% PFA at 4°C for 48 hours, then cryopreserved using 30% sucrose + 0.1% sodium azide in PBS. After a minimum of 48 hours, brains were then embedded in optimal cutting temperature compound (OCT, Fisher Scientific), cryosectioned at 20 µm thickness (Leica, Germany) and transferred to slides for staining. To assess neurogenesis, hippocampal dentate gyrus (DG) sections were stained for Ki67 (proliferative cells), using the primary antibody rabbit anti-Ki67 [LS-C141898; Lifespan Biosciences] and secondary antibody Alexa Fluor 647 goat anti-rabbit [ab150155; Abcam] and doublecortin (DCX) (immature neurons) using the primary antibody guinea pig anti-DCX [AB2253; Millipore], with secondary antibody Alexa Fluor 555 goat anti-guinea pig [A21435; Invitrogen] (**Table 2**). DAPI was used to stain nuclei. Following staining, slides were imaged using confocal microscopy with a 20x objective and a 1.04 µm step size (SP8; Leica), then tiled and analyzed using Imaris software to automatically count the number of Ki67+ and DCX+ cells in the DG region, normalized to the total volume.

#### Whole-mount immunostaining of myenteric neurons

Distal ileum samples from P14 and adult (8-10 week old) mice were placed in cold, oxygenated PBS, opened along the mesenteric border, and gently rinsed in cold PBS to remove luminal contents. For isolation of the muscularis externa, tissues were pinned mucosal side up in a Sylgard-coated dish under a dissecting microscope. The mucosal layer (epithelium and lamina propria) and submucosa (including the submucosal plexus) were carefully removed using fine forceps and micro-scissors, leaving only the longitudinal and circular smooth muscle layers. Isolated muscularis externa was fixed in 4% paraformaldehyde (PFA) for 60 minutes at 4 °C, followed by 3 x 15-min washes in PBS. Tissues were blocked and permeabilized overnight at 4 °C on a shaker in PBS containing 5% normal goat serum (NGS), 10% bovine serum albumin (BSA), and 0.3% Triton X-100. Neurons were stained with ANNA-1 (human anti-HuC/HuD), diluted in blocking buffer and incubated for 96 h at 4 °C with gentle rocking. Tissues were subsequently washed in PBS containing 0.1% Triton X-100 (1 x 5-min wash, followed by 3 x 30 min washes) at 4°C. Secondary antibody was added (goat anti-human IgG [H+L] conjugated to Alexa Fluor 647 diluted in blocking buffer; A21445 [Invitrogen]) and incubated for 24 h at 4 °C and nuclear counterstaining was performed with DAPI (**Table 2**). After washing (3 X 10min in PBS), tissues were mounted serosal side up on glass slides with PBS, and cover-slipped. Slides were stored at 4 °C in the dark until imaging using confocal microscope (Leica SP8).

### Flow Cytometry

Single-cell suspensions of ileum were prepared using a lamina propria dissociation kit (Miltenyi Biotec, Gaithersburg, MD) and gentleMACS tissue dissociator, per manufacturer’s instructions. Briefly, tissue was collected and stored in HBSS with 10 mM HEPES. Samples were cut longitudinally and contents washed off before segmentation into 1 cm pieces. Tissues in HBSS with 10 mM HEPES, 5 mM EDTA, and 5% FBS were placed in shaking incubator at 37°C, 200 rpm for epithelial cell removal. After washing out EDTA from tissues, the samples were incubated at 37°C with MACS digestion enzyme mix before dissociation in the gentleMACS instrument. Cell suspensions were filtered through a 100 μM strainer and washed with FACS buffer for staining.

Cells were stained using a standard staining protocol. After counting single-cell suspension using a hemocytometer with trypan blue exclusion, cells were incubated with Fc block (1:50, anti-CD16-32, 10 μg/mL Tonbo Biosciences, San Diego, CA) for 25 min on ice before addition of antibody cocktail for an additional 30 min (see **Table 2**). After washing with FACS buffer, cells were fixed via incubation with 200 μL BD Cytofix (BD Biosciences, Franklin Lakes, NJ) for 25 min. Samples were run on an LSRII (BD Biosciences, Franklin Lakes, NJ) with BD FACSDiva software (BD Biosciences, Franklin Lakes, NJ). Data was analyzed with FlowJo (Becton Dickinson, Eugene, OR).

### Behavioral testing

#### NOR Task

The NOR task is a two-phase test used to evaluate recognition memory.^8,19,24,26^ For the training phase, mice were transferred to a novel arena (30 cm x 30 cm), containing two identical objects, and allowed to explore for 5 minutes, with each object interaction annotated using automated software (Ethovision, Noldus). Following this phase, mice were returned to their home cage for 45 minutes to recover. For the testing phase, one familiar object from the training phase and one novel object were placed in the arena, and mice were allowed to explore for 5 minutes, with each object interaction scored automatically. An exploration ratio was determined between the number interactions with the novel object over the total number of interactions with both objects. An exploration ratio greater than 50% indicates intact ability to recall familiar objects, with a comparative decrease in exploration ratio indicating a deficit in recognition memory and impaired cognition. Exclusion criteria included an exploration ratio less than 40% or greater than 60% in the training phase, indicating an object preference, or mice with less than 10 total interactions with the objects, indicating lack of engagement during the test.

#### L/D Box

Mice were placed in an arena (40 x 20 cm; 1/3 dark, 2/3 light) for 10 minutes to evaluate anxiety-like behavior.^8,19,24,27^ The time spent in the light compartment was calculated using automated software (Ethovision, Noldus), with greater time spent in the light compartment depicting decreased anxiety-like behavior. Exclusion criteria for this test included mice that made less than 3 transitions between the light and dark compartments and those that indicated an avoidance for the dark box, demonstrated by >500 seconds spent in the light box.

#### OFT

General activity/behavior was tested using the OFT.^8,19,24,26^ Mice were placed in a 30 cm x 30 cm arena and automated software (Ethovision, Noldus) recorded total distance traveled (m) by mice in a 10-minute period to assess general locomotor activity.

### Serum Corticosterone

Blood was collected from neonatal (P14) mice and adult mice (P56) by cardiac puncture and serum was isolated and stored at −80 °C until analysis. Serum corticosterone concentrations were determined using a competitive ELISA kit (Enzo Life Sciences) according to the manufacturer’s small-volume serum protocol. Absorbance was measured at 405 nm with a reference correction at 570 nm using a Synergy H1 plate reader (BioTek Instruments, Inc.). Concentrations were calculated from a four-parameter logistic standard curve and corrected for the sample dilution factor.

### Probiotic muropeptide

#### Isolation of muropeptides

Muropeptides (MP) were isolated from probiotic bacteria according to a previously validated protocol.^28^ Briefly, *Lactobacillus helveticus* (strain R0052) *and L. rhamnosus (*strain R0011) (Lacidofil; kindly provided by Dr. Thomas Tompkins, Lallemand Health Solutions, Canada) was cultured in de Man, Rogosa, and Sharpe (MRS) broth. A starter culture (10 mg/mL in MRS) was incubated overnight at 37°C. The following day, the culture was diluted 1:25 in fresh MRS (20 mL into 500 mL) and grown at 37°C, shaking until OD₆₀₀ ≈ 1.0. Cells were harvested, washed three times in PBS, combined, pelleted (12,000 × g, 10 min), frozen at −80 °C for 24 h, thawed, and refrozen until digestion. Cell pellets were resuspended in sodium dodecyl sulfate, boiled, centrifuged and washed. PGN was weighed and resuspended in lysis buffer (50 mM Tris-HCl pH 8.0, 25 mM NaCl, 2 mM EDTA; 200 µL PBS + 800 µL endotoxin-free water). Mutanolysin (Sigma M4782; 20 U/µL in PBS) was added at 0 h and 24 h (total 6,600 U) while incubated at 37°C, 200 rpm. After 48 h, digests were centrifuged (12,000 × g, 5 min) and the supernatant filtered through 3 kDa Amicon Ultra filters (Millipore UFC500324). Filtrates were washed twice with sterile water, combined, and lyophilized. Muropeptide content was quantified by bicinchoninic acid (BCA) assay (Thermo Fisher Scientific). Preparations were reconstituted in sterile LB broth for oral administration.

#### Analysis of muropeptides

LC/MS was utilized to identify the composition of isolated muropeptides. A Dionex UHPLC coupled to a Fusion Lumos Orbitrap (Thermo Fisher Scientific) was used with C18 column 2.1 × 50 mm (Waters). The LC method was a 0.5 ml min−1 linear gradient starting from 0% A to 50% B in 4 min. Eluent A was 0.1% formic acid in water and Eluent B was 0.1% formic acid in acetonitrile. Thermo Xcalibur Qual Browser was used to process and analyze the data generated. All species showed the expected mass, correct m/z ratio within ± 10 p.p.m. and correct isotopic pattern.

#### *In vitro* splenocyte stimulation

Spleens were aseptically harvested from WT mice and placed in ice-cold PBS. Single-cell suspensions were prepared by gently pressing the tissue through a 100-µm nylon mesh strainer using the plunger of a sterile syringe, followed by rinsing with RPMI-1640 medium supplemented with 10% FBS, 2 mM L-glutamine, and 1% penicillin/streptomycin. The suspension was centrifuged at 1,250 rpm for 5 min at 4°C, and the pellets were lysed using ammonium-chloride-potassium (ACK) lysing buffer. Following lysis, cells were washed and resuspended in RPMI. Cell viability was determined by trypan blue staining, and cells plated at 1.25 × 10^6^ viable cells/well in 24-well plates (500 µL per well) and allowed to equilibrate briefly at 37°C with 5% CO_2_. Cells were then stimulated with one of five treatments: (1) the media alone (control), (2) lipopolysaccharide (LPS; [Invivogen, 2 µg/mL]), (3) muramyl tripeptide-lysine (M-Tri-Lys; [Invivogen, 10μg]), (4) γ-D-glutamyl-meso-DAP (iE-DAP; [Invivogen, 10μg]), or (5) *Lactobacillus* sp. MP; [10μg]). All stimuli were prepared at twice the final concentration in complete medium and added 1:1 (500 µL) to wells, yielding a final culture volume of 1 mL. After 6 h of stimulation, cells were harvested by gentle scraping and centrifugation, and pellets were stored in TRIzol reagent. RNA extraction and cDNA synthesis were performed as described above and gene expression of *Tnf*α and *Il10* was quantified by qPCR.

#### In vivo muropeptide administration

Mice were randomly assigned to one of three groups: Sham+LB, EPEC+LB, or EPEC+muropeptide (EPEC+MP). Muropeptides were freshly reconstituted in LB broth and administered by daily oral administration (5 µL; 10μg) of either muropeptide or LB broth alone (vehicle control) using a pipette tip. The first dose was delivered 1 h prior to EPEC infection (as described above) and repeated every 24 h until P14. Ileal EPEC colonization and inflammation were assessed by histopathology and transcriptomic analysis of inflammatory markers by qPCR at P14.

### Statistics

Results were analyzed by one- or two-way ANOVA with a post hoc Tukey’s or Bonferroni test as appropriate using Prism (GraphPad V10). Data are shown as mean ± SEM and P<0.05 was determined as statistically significant. To determine appropriate sample size, power analysis was conducted (α = 0.05, [1-β] = 0.08) to determine a required n for each test parameter. N represents individual mice, unless otherwise stated.

## RESULTS

### Neonatal EPEC infection disrupts intestinal barrier structure and function at P14

WT and Nod1^ΔIEC^ mice were robustly and reproducibly colonized with EPEC by 7 DPI (P14). Critically, during the acute phase of infection (7 and 14 DPI), fecal pellet bacterial burden was similar in both WT and Nod1^ΔIEC^. In contrast, EPEC clearance was delayed in Nod1^ΔIEC^ mice, with Nod1^ΔIEC^ mice exhibiting significantly higher bacterial loads compared to WT mice at P28 (**Fig 1B**). To assess if neonatal EPEC infection induced structural changes in intestinal architecture and damage, distal ileal tissues were collected at 7 DPI and assessed by histopathology. While no overt histological damage was observed, a reduced crypt-to-villus axis length in WT EPEC-infected mice was seen compared to WT sham mice, consistent with impaired epithelial repair and regeneration (**Fig 1C**). To determine if IEC NOD1 signaling regulated IEC proliferation in healthy controls and during neonatal EPEC infection, we performed confocal microscopy to quantify proliferating Ki67+ IEC (E-cadherin^+^ DAPI^+^) at 7 DPI, the peak of infection. Consistent with the reduced villus-crypt length observed by H&E, confocal microscopy revealed impaired IEC proliferation, indicated by reduced Ki67^+^CDH1^+^DAPI^+^ cells per crypt, in WT EPEC-infected mice compared to WT uninfected mice. EPEC-infected Nod1^ΔIEC^ mice however had equivalent numbers of proliferating IEC (Ki67^+^CDH1^+^DAPI^+^) compared to uninfected Nod1^ΔIEC^ controls (**Fig 1C**). We further assessed if lack of IEC NOD1 altered expression of genes commonly associated with absorbative or secretory lineages of IEC. Increased expression of both *Muc2* (goblet cell marker) and *Slc26a3* (down-regulated in adenoma [DRA]; Cl^-^/HCO^3^ exchanger) mRNA was observed in the distal ileum at P14 in WT EPEC-infected, but not in Nod1^ΔIEC^ mice. (**Fig 1D**). Given the observation of increased Muc2 expression, goblet cells were stained using PAS. Increased total PAS^+^ cells were found in WT EPEC-infected mice, but not Nod1^ΔIEC^ mice (**Fig 1E**). These results demonstrate that NOD1 signaling reduces IEC proliferation and differentiation during enteric bacterial infection with EPEC, including an increase in mucus-producing goblet cells.

### Enteric bacterial infection with EPEC affects the ENS

To assess whether neonatal EPEC infection impacts enteric neurons, expression of neuronal markers was assessed. *Bdnf,* a key neurotropic factor in the GI tract important for regulating physiology,^29^ was significantly increased in WT mice infected with EPEC, but not in Nod1^ΔIEC^ mice (**Fig 2A**). *S100b*, a gene associated with glial activation and neuroimmune signaling, was also increased only in WT mice (**Fig 2A**), together implicating potential early ENS involvement in enteric neuroimmune signaling. Neuronal loss was determined by whole-mount immunostaining of the distal ileum and quantification of neurons in the myenteric plexus. Representative images from WT sham and EPEC-infected mice show a visible reduction in neuronal density following infection, with quantification identifying a significant decrease in neuron counts in WT EPEC-infected mice compared to WT sham-infected controls (**Fig 2B**). The reduction in neuron density was absent in Nod1^ΔIEC^ mice infected with EPEC.

**Figure 2.**
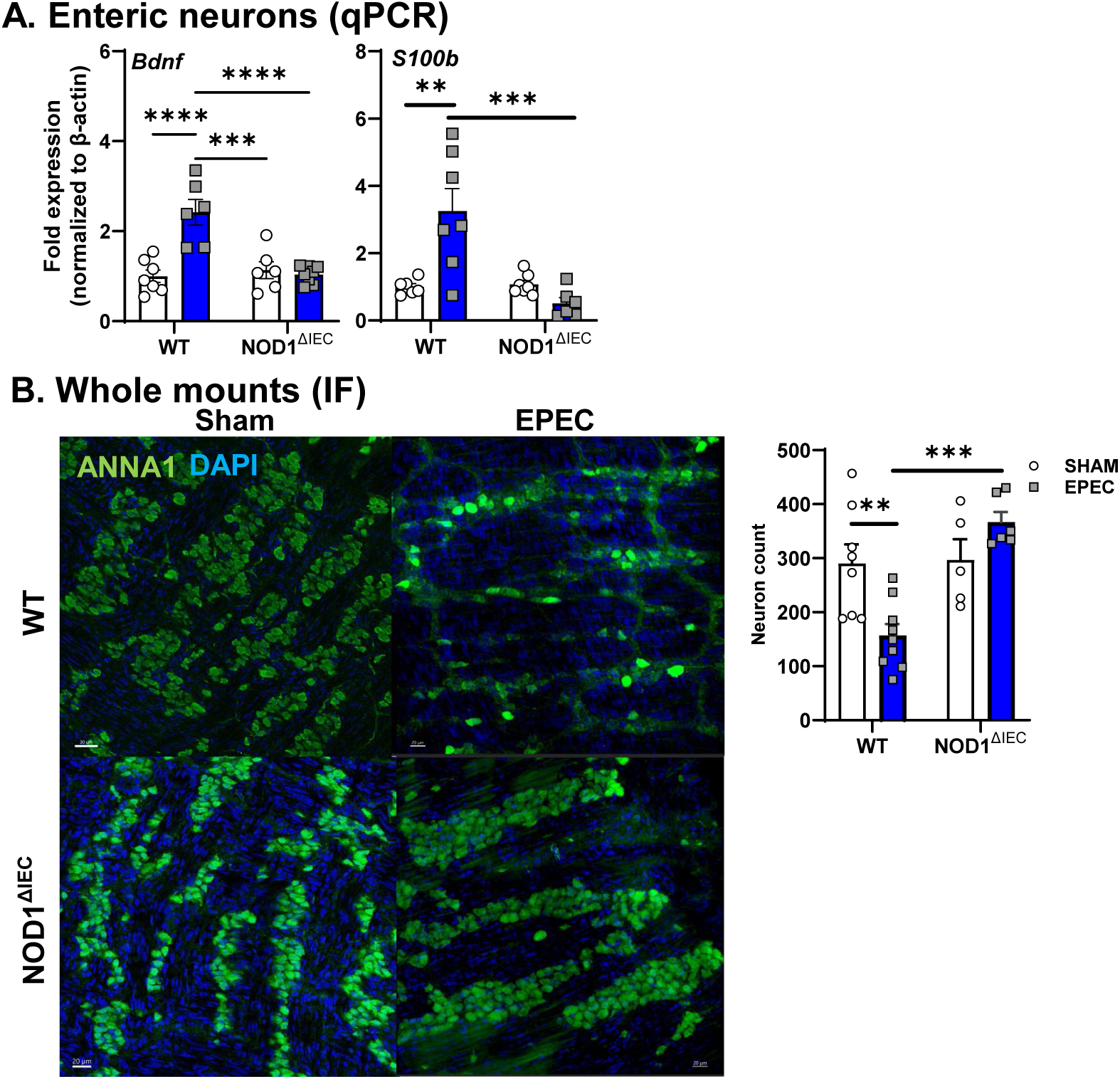
Neonatal EPEC infection disrupts the ENS at P14. Neuronal activation and signaling pathways (*Bdnf*, and *S100b*) in WT and Nod1^ΔIEC^ neonatal mice following EPEC infection (P14) were measured by qPCR (**A**). Whole-mount ileal preparations were stained by ANNA-1 and myenteric neurons counted (**B**). N=6-8 mice per group; mean ± SEM; *P<0.05, **P<0.01, ***P<0.001, ****P<0.001 by 2-way ANOVA,.

### EPEC-induced intestinal inflammation is NOD1 IEC-dependent

Given the significant impact of neonatal infection on physiology in the ileum, early inflammatory response induced by neonatal EPEC were assessed in the distal ileum of WT and Nod1^ΔIEC^ mice. Neonatal EPEC infection elicited a pronounced inflammatory response in WT EPEC-infected mice, with significantly increased expression of pro-inflammatory and host-protective *Il1β*, *Il6*, *Il12*, and *Il22* cytokines compared to WT sham-infected mice (**Fig 3A**). These infection-induced increases in cytokines were not observed in Nod1^ΔIEC^ EPEC-infected mice. Infection in either WT or Nod1^ΔIEC^ mice did not increase expression of *Tnfα* compared to uninfected controls. Expression of prototypical anti-inflammatory cytokine *Il10* was significantly upregulated in WT EPEC-infected mice compared to uninfected control or infected Nod1^ΔIEC^ mice. Conversely, infection did not induce broad expression of regulatory or anti-inflammatory cytokines, as *Tgfb1* was not altered by infection of WT or Nod1^ΔIEC^ mice. Expression of additional PRRs was assessed to better understand whether IEC NOD1 signaling was inducing a proinflammatory state in response to bacterial ligands. Increased expression of *Tlr4* and *Nod2* was observed, indicating amplified innate immune responses in EPEC-infected WT, but not Nod1^ΔIEC^ mice (**Fig 3A**).

**Figure 3.**
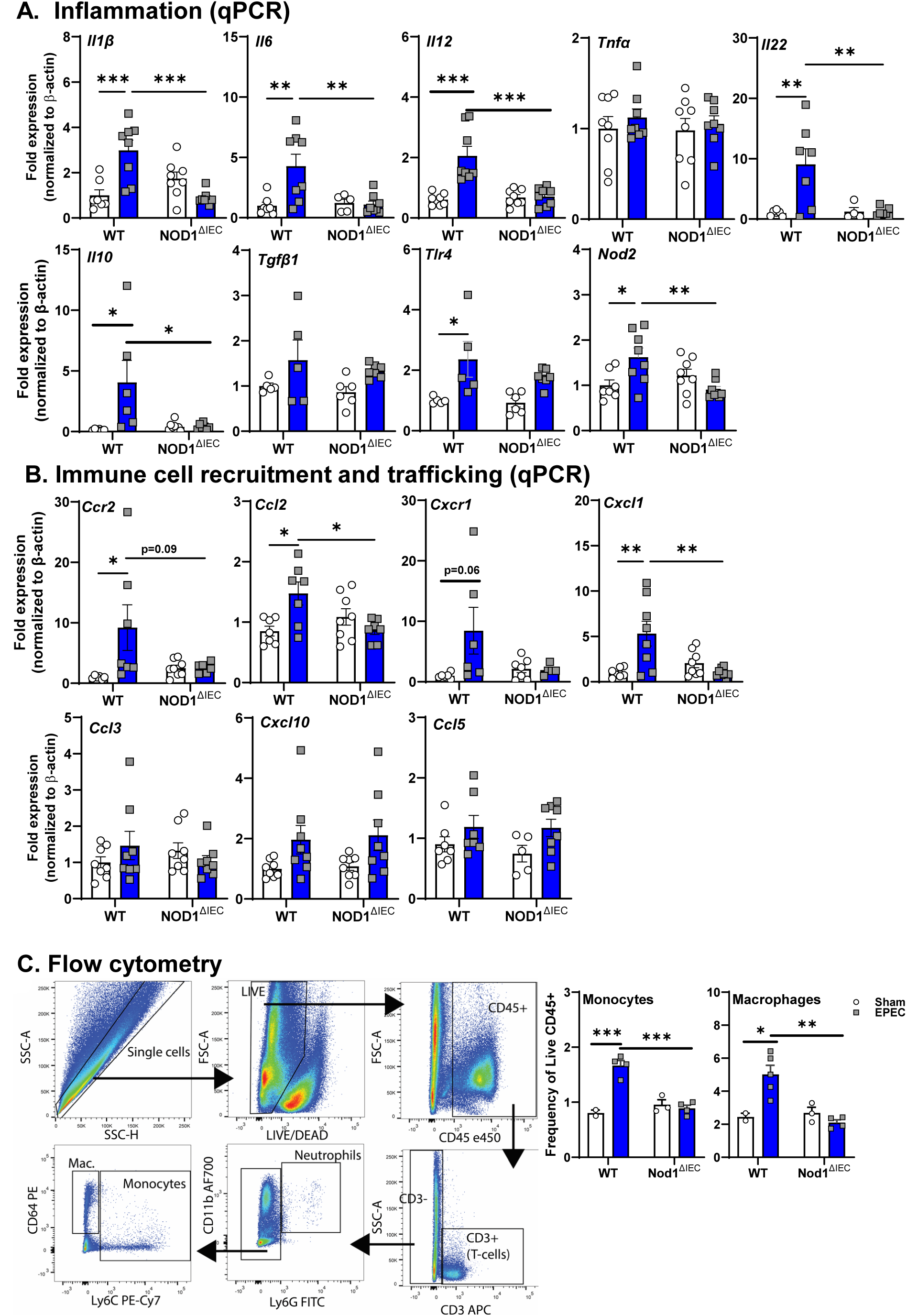
Early life EPEC causes inflammation and alters mucosal immune cell composition in the distal ileum at P14. Inflammatory tone was assessed for pro- inflammatory (*Il1β, Il6, Il12, Tnfα*), regulatory (*Il22, Tgfβ1*) cytokines and pattern recognition receptors (*Tlr2, Nod2*) in WT and Nod1^ΔIEC^ mice following neonatal EPEC infection by qPCR (**A**). Genes involved in immune cell recruitment and trafficking (*Ccr2, Ccl2, Cxcr1, Cxcl1, Ccl3, Cxcl10, Ccl5*) were also quantified by qPCR (**B**). qPCR data are presented as fold change relative to β-actin and normalized to WT sham mice. Representative gating strategy used in flow cytometry to identify live CD45^+^ cells in the distal ileum and distinguish macrophages (CD64^+^ Ly6C^-^, Ly6G^-^), monocytes (CD64^+^, Ly6C^+^), neutrophils (CD11b^+^, Ly6G^+^), and T cells (CD3^+^), with quantification of monocytes and macrophages presented (**C**). Frequencies are expressed as a percentage of live CD45⁺ cells. N=7-8 (qPCR); flow cytometry data is representative of 2 separate experiments (n=7-10 total mice per group); mean ± SEM; *p < 0.05; **p < 0.01; ***p < 0.001 by 2-way ANOVA.

Given the increased expression of select proinflammatory cytokines, we assessed if chemokine expression and recruitment of specific immune cells occurred during neonatal EPEC infection of the ileum, and whether these are dependent on NOD1 signaling in IEC. Infection in WT mice resulted in significantly increased expression of *Ccr2* and its cognate ligand *Ccl2* (**Fig 3B**). Similarly, *Cxcr1* was also elevated (p=0.06) along with its cognate ligand *Cxcl1* (**Fig 3B**). This infection induced increase was dependent on NOD1 signaling in IEC as EPEC-infection did not increase expression of these genes in Nod1^ΔIEC^ mice. Infection did not increase the expression of all chemokines and receptors, as *Ccl3*, *Ccl5*, and *Cxcl10*, were not impacted in WT EPEC-infected mice (**Fig 3B**).

With these changes in key chemokines that could alter monocyte homing during infection, we characterized immune cell recruitment in the distal ileum following neonatal EPEC infection using flow cytometry (**Fig 3C**). EPEC infection of neonatal mice significantly increased the frequency of monocytes (CD64⁺^/-^Ly6C⁺) in WT ileum compared to WT sham-infected mice, while there was no change in monocytes in uninfected or infected Nod1^ΔIEC^ mice. Similarly, macrophage (CD64⁺Ly6C⁻) frequency was significantly elevated in WT EPEC-infected mice compared to WT sham mice and Nod1^ΔIEC^ EPEC-infected mice (**Fig 3C**). No significant differences were observed in T-cells or neutrophils (data not shown). These findings demonstrate that EPEC infection drives the recruitment of specific immune cell populations in the neonatal small intestine in a Nod1-dependent manner.

### Neonatal EPEC infection leads to persistent disrupted intestinal barrier function and EPEC-induced GI inflammation persists into adulthood in WT mice

The long-term consequence of neonatal EPEC infection was assessed in adult mice (P56) after pathogen clearance. Physiology of the ileum including, baseline ion secretion, and permeability was assessed in Ussing chamber studies. Ion secretion, as indicated by Isc, was not significantly different in either WT or Nod1^ΔIEC^ mice that were sham- or EPEC-infected as neonates. Prior neonatal infection, however, significantly increased ileal permeability as measured by G and FITC flux in WT, but not Nod1^ΔIEC^ mice (**Fig 4A**). Despite the lack of effect on ion secretion, increased expression of the chloride-sulfate anion transporter *Slc26a3* (DRA) was observed in adult mice previously infected with EPEC. Expression of *Muc2* and total number of PAS^+^ goblet cells was not different in adult WT or Nod1^ΔIEC^ mice irrespective of prior infection status suggesting the presence of an intact mucus barrier (**Fig 4B**). Similarly, the villus-crypt axis and the total number of Ki67^+^ IEC were not significantly impacted by neonatal EPEC infection in either WT or Nod1^ΔIEC^ mice (**Fig 4C**), suggesting that EPEC-induced deficits in IEC kinetics were transient, and limited to the period of active infection.

**Figure 4.**
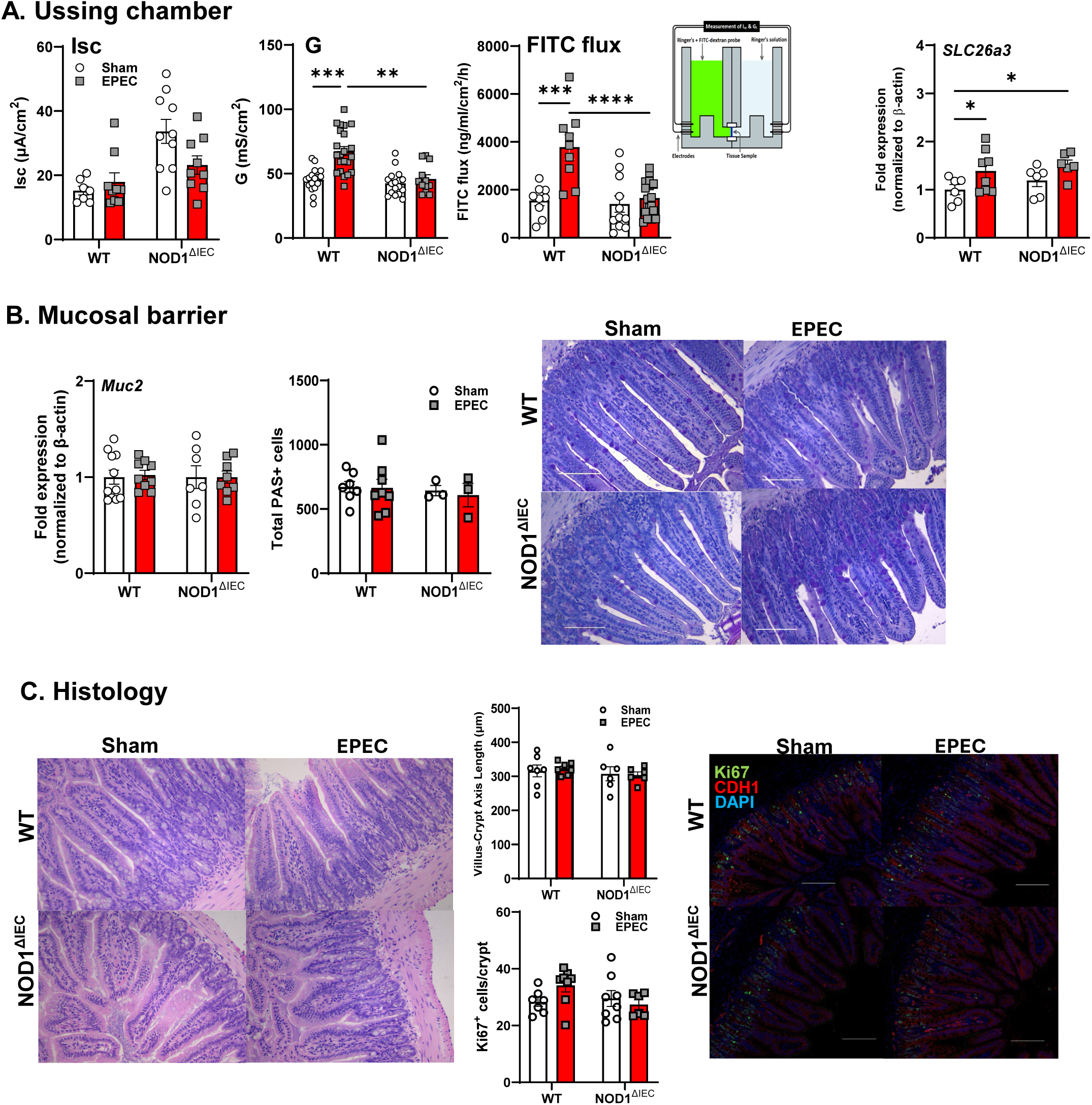
Neonatal EPEC infection leads to persistent disrupted intestinal barrier function in adult mice, which is absent in Nod1^ΔIEC^ mice. Ussing chamber studies assessed short-circuit current (Isc), transepithelial conductance (G), and FITC-dextran flux in ileum from adult WT and Nod1^ΔIEC^ mice either sham- or EPEC-infected neonatally (**A**). Epithelial barrier markers (Slc26a3) were assessed by qPCR (**A**). Goblet cells were assessed by qPCR and PAS staining to quantify Muc2 expression and quantify goblet cell numbers (**B**). Histology was used to assess the villus-crypt axis (length) and quantify the number of proliferative IEC (Ki67+) (**C**). N=8-12; mean ± SEM; *p < 0.05; **p < 0.01; ****p < 0.0001 by 2-way ANOVA.

As intestinal barrier function is a critical determinant of health, we assessed if these persistent barrier defects in adult mice infected as neonates lead to expression of inflammatory cytokines. Adult WT, but not Nod1^ΔIEC^, mice previously infected as neonates had increased expression of *Il6*, *Il12*, *Ifnγ, and Il22* (**Fig 5A**). In contrast, *Il1β* and *Tnfα* were not significantly different across any of the experimental groups. Persistently increased expression of *Nod2* was found in WT EPEC compared to WT sham and Nod1^ΔIEC^ EPEC groups, but no changes were found in expression of the glial marker S100b (**Fig 5A**). Lastly, analysis of chemokine and receptor expression revealed persistently increased *Ccl2* and *Cxcr1* in adult WT, but not Nod1^ΔIEC^, mice previously infected as neonates, with no changes in *Cxcl1*, or *Ccr2* expression (**Fig 5A**). Finally, myenteric neurons were quantified and in contrast to P14 mice, no changes in total numbers were seen in either WT or Nod1^ΔIEC^ mice neonatally infected with EPEC, suggesting that the number of enteric neurons recovers post-infection (**Fig 5B**).

**Figure 5.**
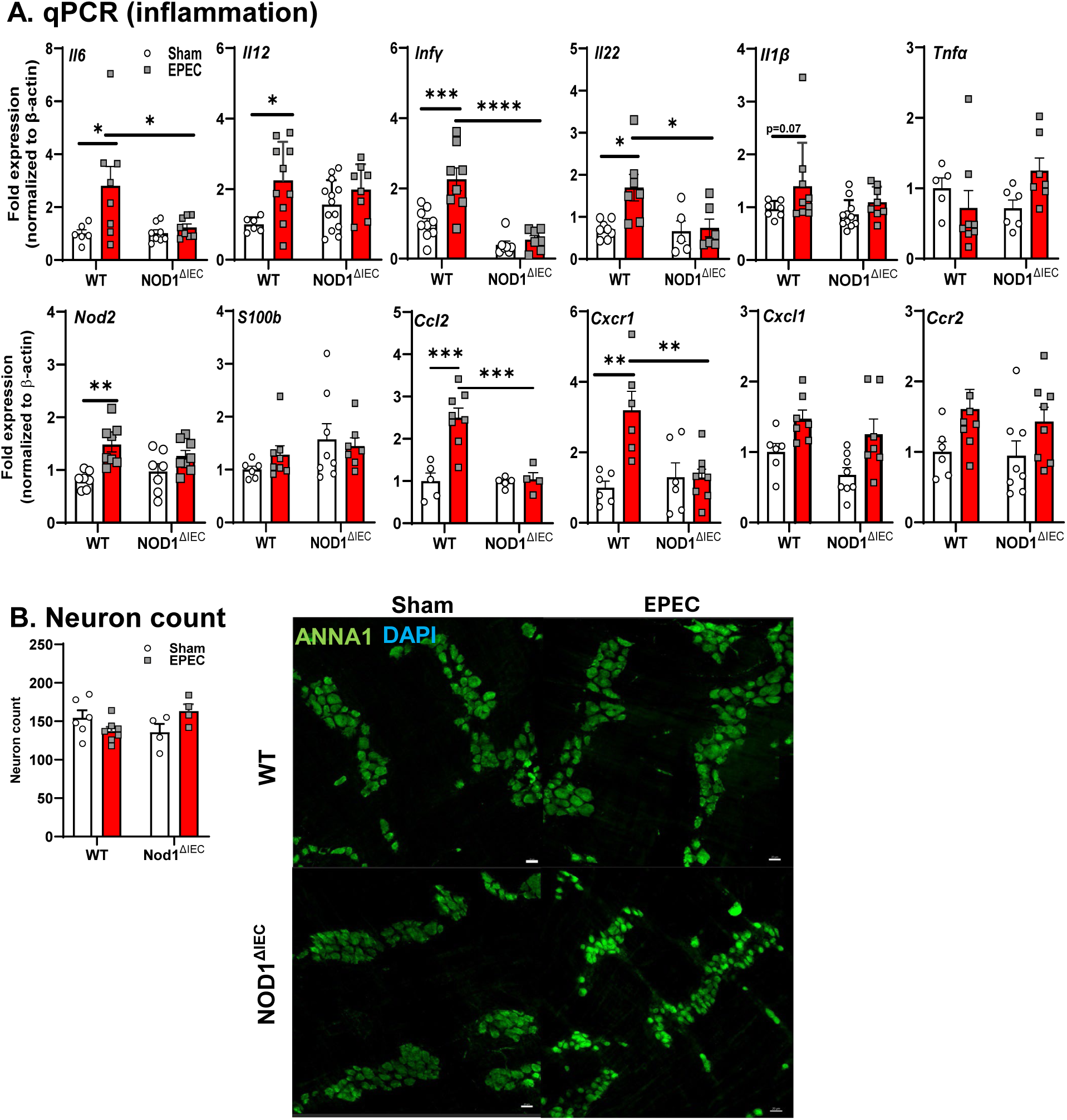
**Neonatal EPEC infection causes persistent ileal inflammation without impacting the ENS in WT mice**. Cytokines (*Il6, Il12, Ifnγ, Il22, Il1β,* and *Tnfα*), *Nod2, S100b*, and chemokines (*Ccl2, Cxcr1, Cxcl1* and *Ccr2*) were measured by qPCR in distal ileum from WT and Nod1^ΔIEC^ mice (**A**). Myenteric neurons were stained (ANNA-1) and quantified in WT and Nod1^ΔIEC^ mice (**B**). N=8-12; mean ± SEM; *p < 0.05; **p < 0.01; ****p < 0.0001 by 2-way ANOVA.

### Neonatal infection causes cognitive deficits, neuroinflammation, altered hippocampal neurogenesis, and neuroplasticity in adulthood

To assess the role of IEC NOD1 on the long-term neurobehavioral impact of neonatal EPEC infection, we performed behavioral testing in adult mice starting at P56. Recognition memory, as determined using the NOR task, was impaired in neonatally EPEC-infected WT mice, but not Nod1^ΔIEC^ mice, as determined by a significant reduction in exploration ratio compared to WT sham mice (**Fig 6A**). In contrast, there were no significant differences in anxiety-like behavior as measured by time spent in the light compartment of the L/D box, nor in locomotor activity assessed by total distance traveled in the OFT across any group (**Fig 6A**).

**Figure 6.**
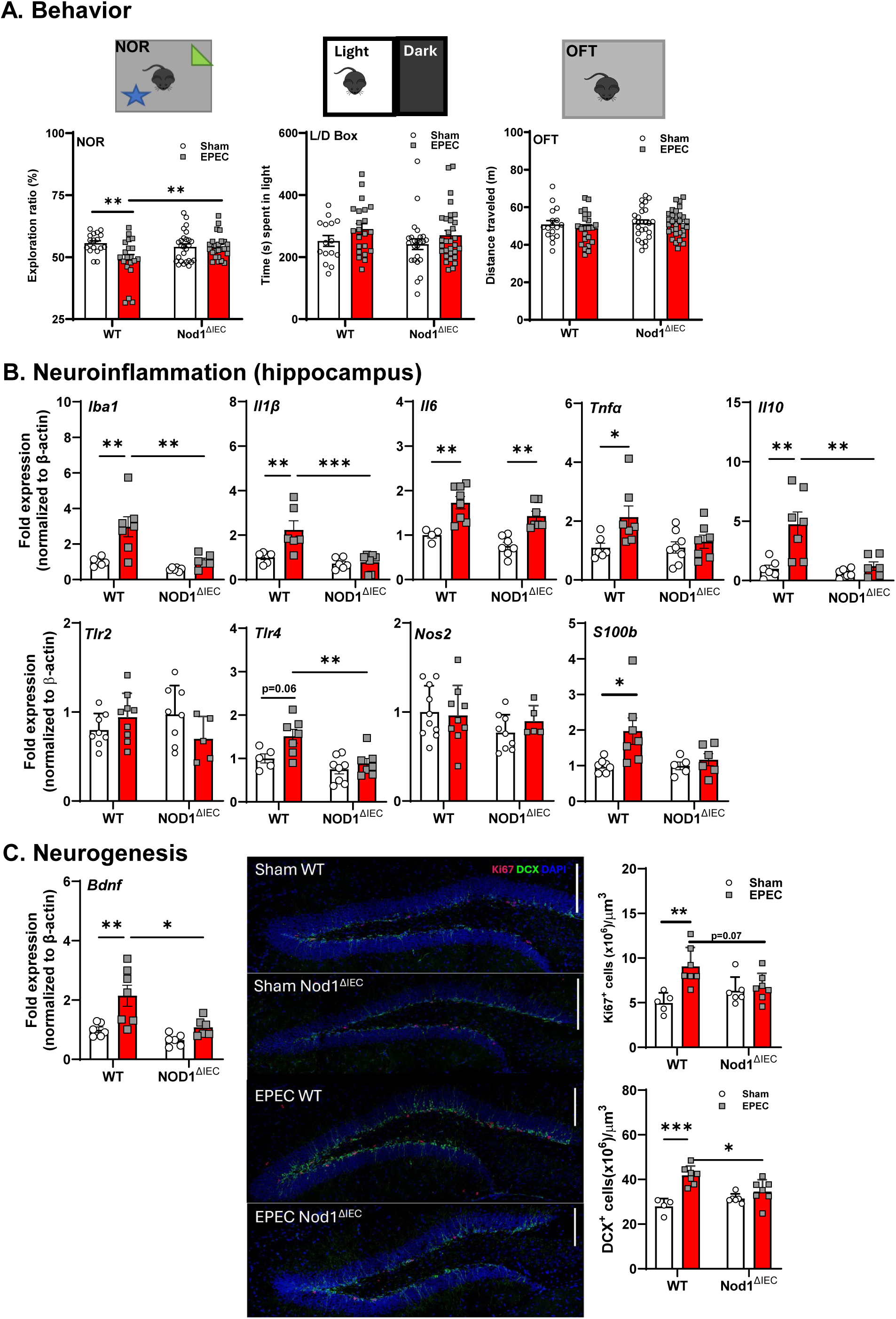
Early-life enteric infection leads to behavioral deficits coupled with neuroinflammation and decreased neurogenesis in adulthood. Behavior was assessed in adult mice using the novel object recognition (NOR) task, light/dark (L/D) box, and open field test (OFT) to evaluate cognitive, anxiety-like, and locomotor behaviors in WT and Nod1^ΔIEC^ mice (**A**). mRNA expression of inflammatory and glial activation markers in the hippocampus of adult mice following neonatal EPEC infection was performed, including Iba1, *Il1β*, *Il6*, *Tnfα*, *Il10*, *Tlr2*, *Tlr4, Nos2*, and *S100b* by qPCR (**B**). Data are presented as fold change relative to β-actin and normalized to WT sham mice. To assess neurogenesis, fold-change in hippocampal *Bdnf* mRNA expression was performed along with immunofluorescence of Ki67⁺ (proliferation) and doublecortin (DCX⁺ - new neurons) positive neurons in the dentate gyrus in WT and Nod1^ΔIEC^ mice at P56 (**C**). Representative immunofluorescence images show Ki67 (red), DCX (green), and DAPI (blue) staining. Scale bars = 100 μm. Behavior n=15-28, qPCR n=5-8, IF n=5-7; mean ± SEM; *p < 0.05; **p < 0.01; and ****p < 0.0001 by 2-way ANOVA.

Our previous studies found that neuroinflammation was not observed in neonatal mice following EPEC infection (P21), but rather developed in adulthood, suggesting a delayed response within the CNS to enteric infection.^8^ Here, we confirmed those observations in adulthood, with increased hippocampal *Iba1* expression, a marker of microglia, in WT mice infected with EPEC as neonates compared to non-infected controls and Nod1^ΔIEC^ mice. In addition, *Il1β*, *Il6*, and *Tnfα* were significantly upregulated in adult WT mice previously infected with EPEC as neonates compared to WT sham mice and Nod1^ΔIEC^ mice (**Fig 6B**). *Il10* was also significantly increased in WT EPEC-infected mice compared to sham mice, potentially indicating a compensatory anti-inflammatory response that may be driven by heightened pro-inflammatory signaling in WT mice (**Fig 6B**). With respect to PRR expression, *Tlr2* remained unchanged whereas *Tlr4* expression was significantly increased in WT EPEC-infected mice compared to both WT sham-infected controls and Nod1^ΔIEC^ mice (**Fig 6B**). *S100b* showed modest but significant elevation (P<0.05) in WT mice previously infected with EPEC compared to WT sham mice, suggesting the activation of glial cells (**Fig 6B**).

Given the evidence of hippocampal neuroinflammation in adult WT mice neonatally infected with EPEC, neurogenesis was assessed. Adult WT mice neonatally infected with EPEC had a significant increase in hippocampal *Bdnf* expression compared to WT sham mice, as seen in our prior studies (**Fig 6C**).^8^ However, this upregulation was significantly blunted in Nod1^ΔIEC^ EPEC-infected mice, with gene expression levels comparable to sham-infected mice. These findings point to lasting deficits in hippocampal plasticity following neonatal infection, dependent in part on intestinal epithelial NOD1 signaling. To further evaluate long-term the effects of neonatal EPEC infection on hippocampal neurogenesis, we quantified proliferating (Ki67⁺) and immature neuronal (DCX⁺) cells in the dentate gyrus region of the hippocampus of adult mice (P56). WT EPEC-infected mice, but not Nod1^ΔIEC^ EPEC-infected mice, showed a significant reduction in Ki67⁺ cell density compared to WT sham mice (**Fig 6C**). Similarly, DCX⁺ neuroblast density was significantly increased in WT EPEC-infected mice relative to sham-infected, but not in Nod1^ΔIEC^ EPEC-infected mice (**Fig 6C**).

### Early-life EPEC infection results in sustained systemic HPA axis activation and altered serotonergic signaling in WT mice

5-HT has a critical function in maintenance of gut-brain communication, therefore we assessed expression of genes critical for serotonergic signaling in the neonatal ileum and in adult ileum and brain following infection. In neonatal mice, *Tph2, and Sert* expression in the ileum was significantly elevated in EPEC-infected WT, but not Nod1^ΔIEC^ or uninfected mice.

In contrast, *Tph1* showed no significant changes at 7 DPI (P14), suggesting that 5-HT production by enterochromaffin cells was not impacted by infection (**Fig 7A**). Increased expression of the 5-HT receptors *5htr2c* and *5htr4* were also observed only in the infected WT mice.

**Figure 7.**
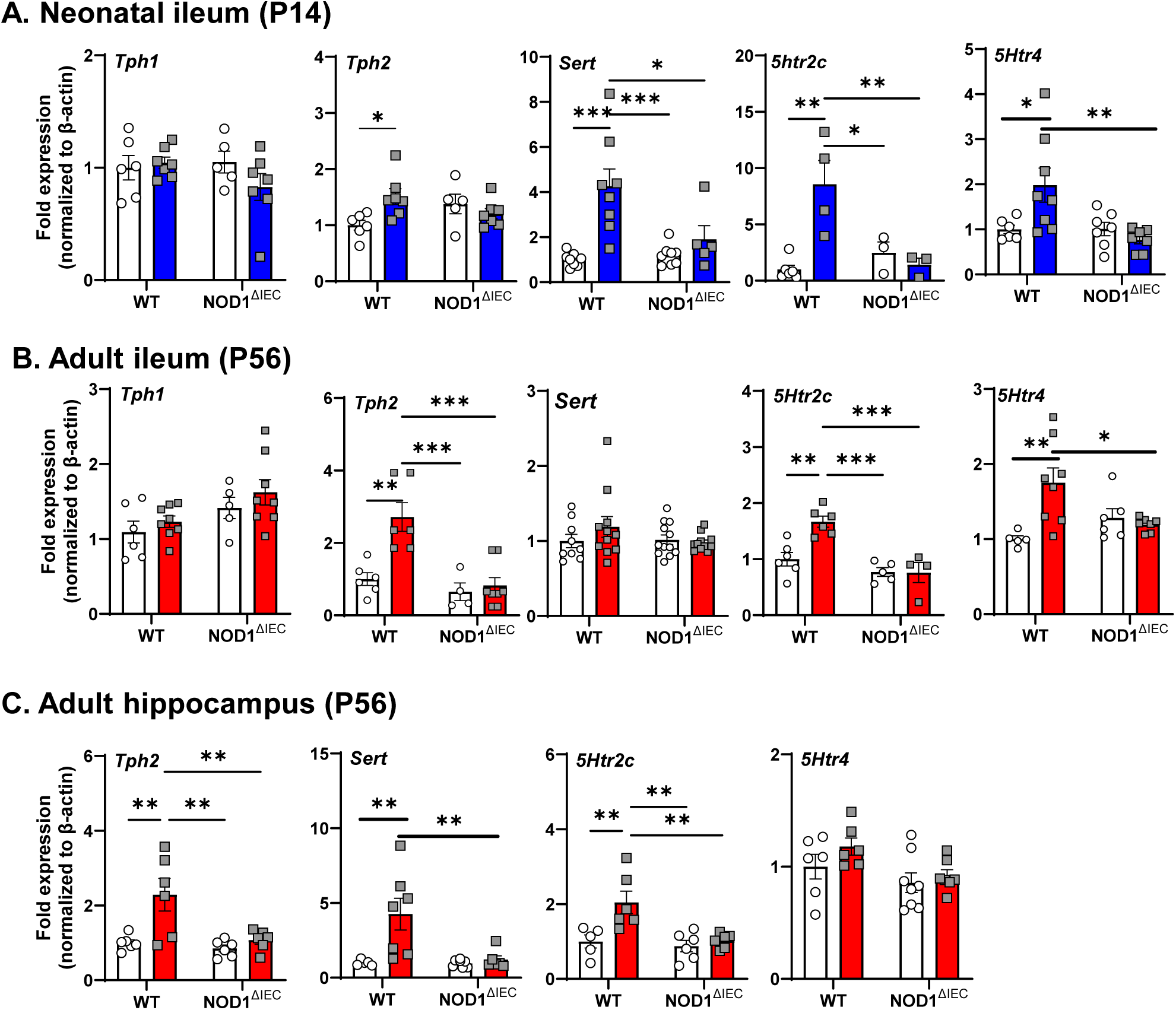
Neonatal EPEC infection impacted serotonin signaling in both the gut and brain via IEC NOD1. Expression of serotonin (5-HT)-related genes including *Tph1*, *Tph2*, *Sert*, and *5htr2c* at both P14 (**A**) and adult (**B**) ileum and in the adult hippocampus was determined by qPCR (**C**). N=5-8; mean ± SEM; *p < 0.05; **p < 0.01; and ****p < 0.0001 by 2-way ANOVA.

By adulthood (P56), EPEC-infected WT mice similarly showed increased expression of *Tph2, 5htr2c,* and *5Htr4* compared to WT sham mice in distal ileum (**Fig 7B**), suggesting prolonged enteric serotonergic dysregulation in the ileum caused by neonatal bacterial infection. These impairments in 5-HT signaling were absent in Nod1^ΔIEC^ mice. In the adult hippocampus, WT EPEC-infected mice also demonstrated significant upregulation of *Tph2*, *Sert,* and *5htr2c,* but not *5Htr4,* compared to sham-infected controls, which were again absent in Nod1^ΔIEC^ mice (**Fig 7C**). Together these findings highlight that neonatal EPEC infection persistently disrupts 5-HT signaling in both the gut and brain.

Given the interaction between 5-HT and HPA axis activation, stress response-related gene expression was assessed. Neonatal EPEC infection induced persistent alterations in host stress-related signaling along the MGB axis. In the neonatal distal ileum (P14), EPEC infection markedly increased expression of the stress receptor gene *Crfr2* in WT EPEC-infected mice compared to sham-treated mice, an effect that persisted into adulthood, but was absent in Nod1^ΔIEC^ mice **(Fig 8A)**. On the other hand, *Gr* expression was unchanged in both neonatal and adult ileum. In the adult hippocampal tissue from WT EPEC-infected mice, elevated *Crfr2* and *Nos2* expression was observed vs. WT sham controls, and increased *Gr* expression compared to all other groups (**Fig 8B**). This increased GR expression may be associated with disrupted HPA axis feedback inhibition. These hippocampal changes were absent in Nod1^ΔIEC^ mice. Functionally these gene expression changes led to increased serum corticosterone levels, which were elevated in WT mice previously infected with EPEC-infected mice, but only in adulthood (**Fig 8C**).

**Figure 8.**
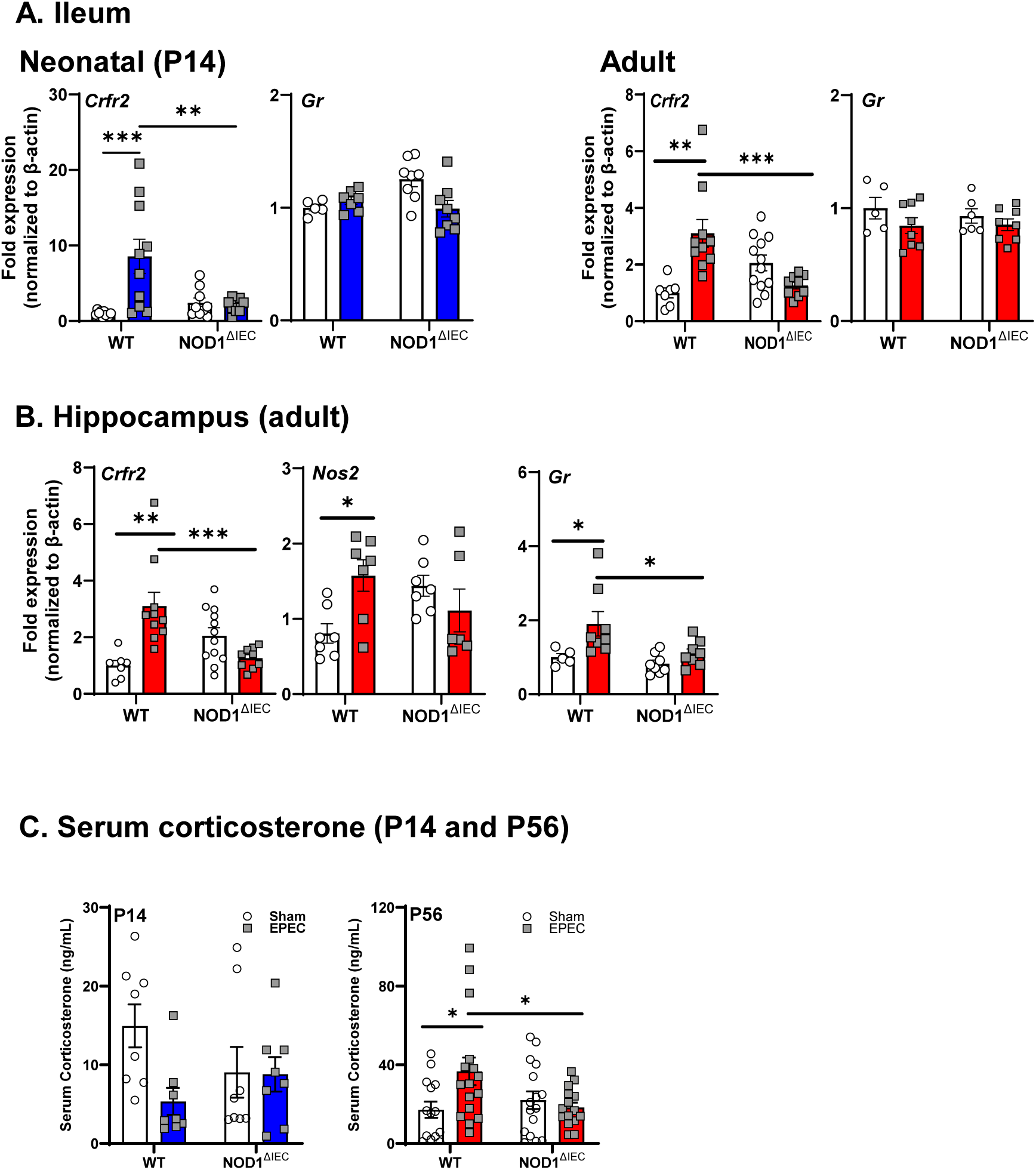
Early-life EPEC infection results in sustained systemic HPA axis activation. HPA axis signaling markers (*Crfr2, Gr*) were measured in the neonatal (P14) and adult ileum (**A**), and adult hippocampus (**B**) following neonatal EPEC infection in WT and Nod1^ΔIEC^ mice by qPCR. Serum corticosterone was assessed in both neonatal (P14) and adult (P56) WT and Nod1^ΔIEC^ mice (**B**). qPCR n=5-6, CORT EIA n=8-16; mean ± SEM; \**p* < 0.05*, **p < 0.01, ***p < 0.001* by 2-way ANOVA.

### Probiotic muropeptide supplementation mitigates EPEC-induced neonatal ileal inflammation

To determine whether probiotic muropeptides could be beneficial following neonatal EPEC infection, they were isolated from probiotic PGN and analyzed by LC/MS. Analysis revealed a complex mixture of monomeric, dimeric, and trimeric muropeptides isolated from probiotic Lactobacillus strains (**Fig 9A**). Relative quantification, based on total ion counts, indicated dominance of crosslinked structures, with Tetra-N-Tetra-N trimer being most abundant (32.94% of total ion signal), followed by Tetra-N-Tetra-N dimer (21.56%) and Tri-N-Tetra-N trimer (16.94%). Monomeric species, including Tri and Tetra forms, were detected at trace levels (<0.3% each). The structural analysis confirmed the presence of N-acetyl-glucosamine (GlcNAc)–N-acetyl-muramic acid (MurNAc) disaccharide units linked to canonical stem peptides (L-Ala–D-Gln–L-Lys–D-Ala), bridged by peptide crosslinks characteristic of Gram-positive PGN.

**Figure 9.**
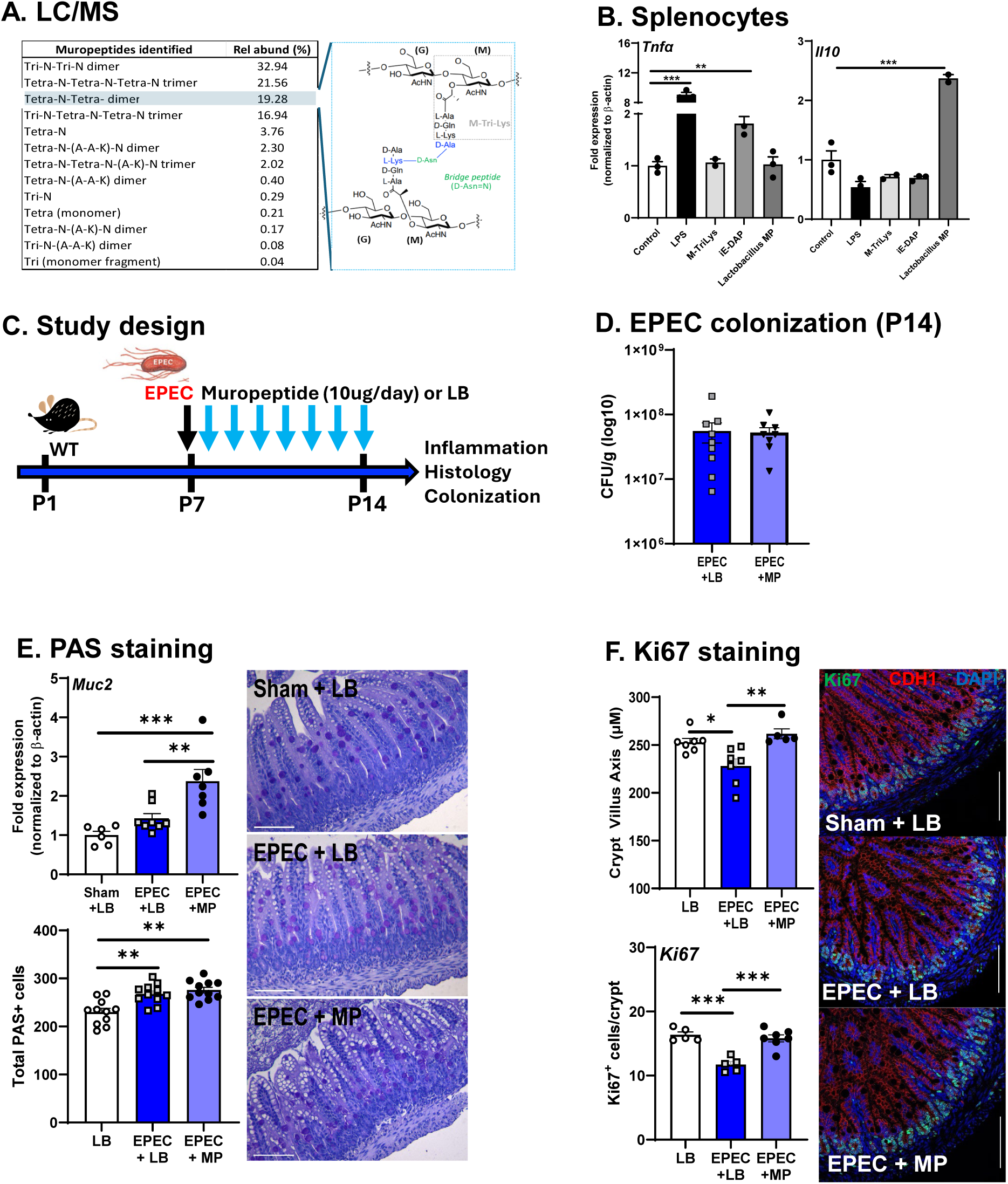
Probiotic muropeptide supplementation mitigates EPEC-induced neonatal epithelial cell deficits. Peptidoglycan from probiotic strains (*L. helveticus/L. rhamnosus*) was isolated and characterized by LC/MS **(A)**. Murine splenocytes were cultured *in vitro* and stimulated with bacterial ligands (LPS, M-Ti-Lys, IE-DAP, and Lactobacillus muropeptide [MP]) and *Tnfα* and *Il10* measured by qPCR. Probiotic muropeptides were administered *in vivo* in neonatal WT mice (**C**). Bacterial burden in the distal intestine (log10 CFU/g feces) of neonatal mice orally infected with EPEC and supplemented daily from P7 to P14 with either vehicle (LB) or bacterial muropeptides was quantified (**D**). Expression of *Muc2* was measured by qPCR and goblet cell were stained using PAS and positive cells quantified (**E**). Epithelial cell proliferation was assessed by measuring the villus-crypt axis and staining with Ki67 to quantify positive cells in the distal ileum (**F;** Ki67 [green], CDH1 [red], DAPI [blue]). Splenocytes n=2-3, colonization n=8-9, IF n=6-10; mean ± SEM; *p < 0.05, **p < 0.01, ***p < 0.001.

Using an *in vitro* assay to assess immunogenicity, isolated murine splenocytes were stimulated with probiotic MP, and cytokine expression (*Tnfα* and *Il10*) was measured. As expected, *Tnfα* expression was significantly increased by LPS or iE-DAP stimulation, whereas M-Tri-Lys and probiotic MP stimulation did not increase Tnfα expression relative to control (**Fig 9B**). In addition, probiotic MP markedly upregulated *Il10* response from splenocytes compared to all other groups (**Fig 9B**). These data indicate that the probiotic-derived muropeptide fraction is pro-regulatory in splenocytes, potentially capable of eliciting an anti-inflammatory response.

To assess whether probiotic muropeptides could ameliorate EPEC-induced GI inflammation, mice were orally supplemented daily for 7 days (**Fig 9C**). To determine whether supplementation with probiotic muropeptides, including the beneficial M-Tri-Lys, alters colonization during neonatal EPEC infection, bacterial burden was quantified in fecal samples at P14. No significant differences in EPEC colonization were observed between the EPEC+LB and EPEC+MP treated mice (**Fig 9D**). This indicates that muropeptide supplementation did not impact colonization or limit acute infection in EPEC-infected mice. Given the important role for the mucus layer in maintaining epithelial barrier integrity, we assessed the impact of EPEC+MP on mucins and goblet cells. Expression of *Muc2* was increased in both EPEC+LB treated and EPEC+MP treated mice compared to vehicle controls, which was coupled with a significant increase in total PAS-positive cells compared with uninfected LB controls, findings consistent with mucosal remodeling and goblet cell hyperplasia during infection (**Fig 9E**). Notably, daily oral supplementation with muropeptides did not prevent EPEC-induced increases in PAS^+^ goblet cells compared to EPEC-infected mice receiving LB (vehicle). These findings indicate that, while muropeptide administration did not prevent EPEC-induced goblet cell expansion, this feature may serve as a protective effect on epithelial homeostasis. Lastly, epithelial cell kinetics were assessed histologically. EPEC-induced decreases in crypt/villus axis were prevented by muropeptide supplementation, which were coupled to restoration in numbers of Ki67^+^ epithelial cells (**Fig 9F**). This supports the idea that muropeptides enhance mucus layer integrity, a critical barrier against luminal pathogens, thereby reinforcing epithelial defenses in early life.

To evaluate the immunomodulatory potential of muropeptides during early-life EPEC infection, mRNA expression of key pro- and anti-inflammatory mediators was assessed in the distal ileum at P14. As expected, EPEC+LB supplementation increased *Il1β*, *Il6*, *Il12, Il22* and *Tnfα* compared to both vehicle control mice, which was prevented in EPEC+MP treated mice (**Fig 10A**). With respect to regulatory cytokines, *Il10* was found to be elevated in EPEC+LB mice compared to vehicle control, but not in EPEC+MP mice (**Fig 10A**). No differences were observed between vehicle control and EPEC+MP mice, indicating that the supplementation did not broadly dampen anti-inflammatory responses, but rather rebalanced the inflammatory environment. The increased expression of the glial marker *S100b* in EPEC+LB mice was also inhibited in EPEC+MP treated mice (**Fig 10A**). PRR receptors were also affected by muropeptide supplementation, with EPEC+LB supplementation causing upregulated *Tlr2*, *Tlr4*, and *Nod2* compared to vehicle control, which was prevented in EPEC+MP-treated mice (**Fig 10B**). This downregulation of TLR-driven microbial sensing may help limit overactivation of innate immune responses caused by EPEC-infection.

**Figure 10.**
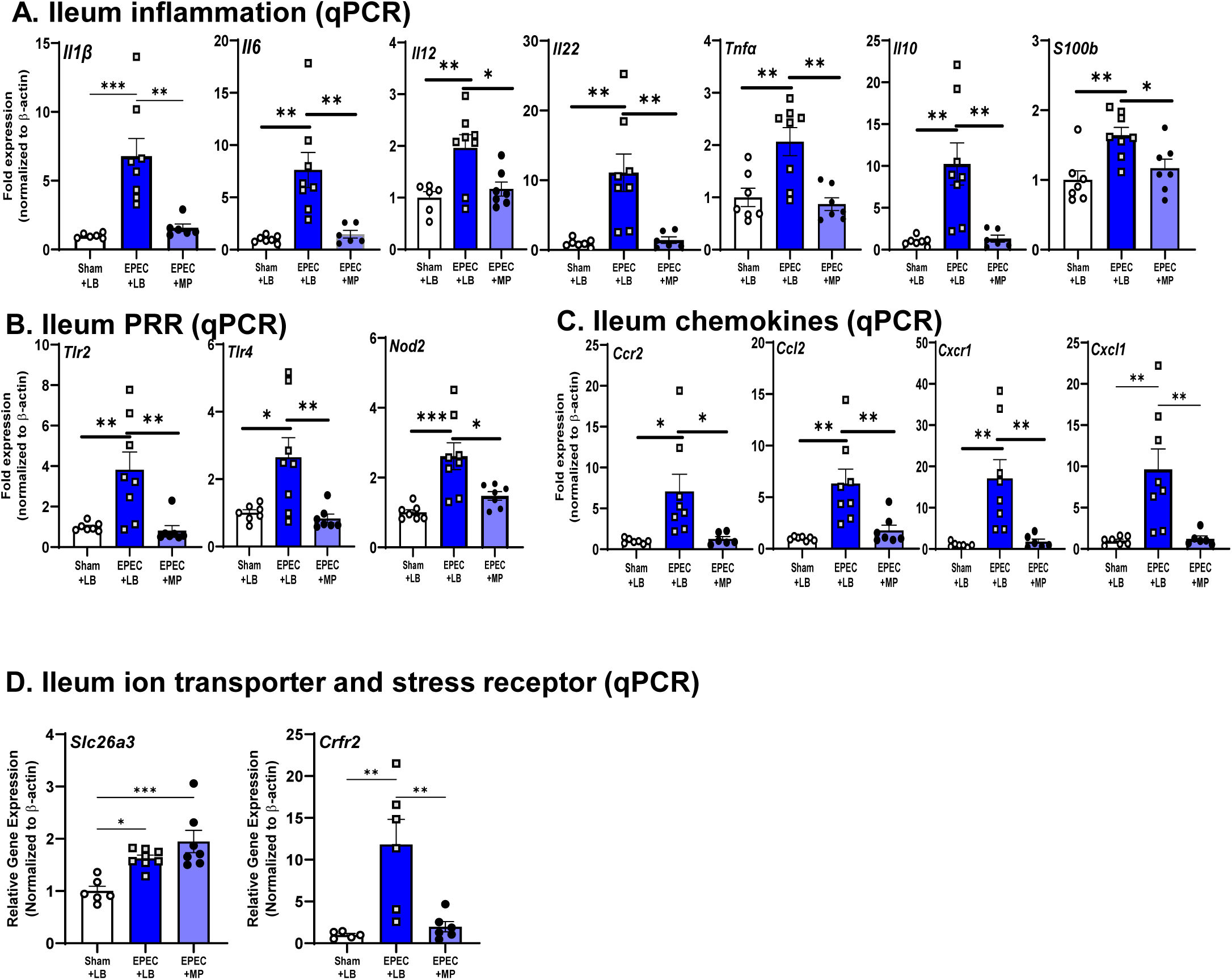
Immune responses elicited by neonatal EPEC infection were prevented by probiotic muropeptide supplementation. mRNA expression of pro-inflammatory cytokines (*Il22*, *Il6*, *Il12*, *Tnfα*, *Il1β*), anti-inflammatory/regulatory cytokines (*Il10*), and the glial cell marker S100b was assessed in the ileum (**A**). Pattern recognition receptors (*Tlr2*, *Tlr4*, *Nod2*) (**B**), and chemokines (*Ccl2*, *Cxcl1*, *Ccr2*, *Cxcr1*) (**C**) were also quantified in the distal ileum at P14 by qPCR. Lastly, the epithelial transporter (*Slc26a*) and stress receptor (*Crfr2*) were quantified by qPCR (**D**). N=5-8; mean ± SEM; *p<0.05, **p<0.01, ***p<0.001 by one-way ANOVA.

Chemokine profiles revealed similar trends. EPEC+LB mice displayed higher expression of *Ccr2, Ccl2*, *Cxcr1* and *Cxcl1* compared to vehicle controls, which was prevented by MP supplementation. These findings indicate that muropeptides may limit chemokine-driven recruitment of monocytes and neutrophils to the infected mucosa (**Fig 10C**). Increased expression of *Slc26a3* in EPEC-infected mice was not improved by MP treatment, but EPEC-induced elevated expression of *Crfr2* was similar to sham control levels in the EPEC+MP group (**Fig 10D**). The normalization of *Crfr2* expression with muropeptide supplementation suggests a potential mechanism by which these bioactive molecules may modulate GI-derived neuroendocrine signaling. This raises the possibility that muropeptides could mitigate the downstream brain and behavioral alterations we observed in EPEC+LB mice.

Lastly, given the EPEC-induced impairments seen in the myenteric plexus at P14, as determined by significant decrease in total neuron numbers, quantification of enteric neurons was performed. Excitingly, supplementation with muropeptide restored neuron counts in EPEC+MP treated mice to similar numbers seen in sham (LB) treated mice (LB: 371 ± 15 vs. EPEC+MP: 367 ± 23 neurons; n=4 per group, P = NS), highlighting the ability of MP to protect myenteric neurons following bacterial infection.

Collectively, treatment with probiotic-derived muropeptides blunts EPEC-induced pro-inflammatory cytokine and chemokine cascades, downregulates excessive PRR-mediated microbial sensing, normalizes innate immune receptor expression, preserves functional mucus barrier integrity and restores the ENS. These converging effects suggest a protective mechanism likely involving Nod2-dependent pathways, positioning muropeptides as promising candidates for mitigating infection-induced inflammation in the immature gut.

## DISCUSSION

Neonatal enteric infections remain a critical global health concern in part due to their potential long-term effects on GI physiology, immunity, and neurodevelopment.^5,30^ Previously, we showed that early-life EPEC infection causes acute neonatal GI inflammation leading to disruption of intestinal immune homeostasis and mucosal barrier function, coupled with CNS deficits (neuroinflammation and impaired neurogenesis), and persistent behavioral changes in adulthood, disrupting the MGB axis.^8^ Here, we extend these findings by demonstrating that intestinal epithelial NOD1 signaling not only regulates local immune responses but also shapes long-term neuroimmune outcomes following enteric infection, and that supplementation with probiotic muropeptide can mitigate infection-induced MGB axis deficits.

Innate immune responses are important for clearance of enteric bacterial pathogens.^31^ EPEC robustly and reproducibly infects and colonizes neonatal mice, with clearance occurring between 14- and 21-days post-infection in WT mice.^8,11^ However, in the absence of IEC NOD1, EPEC clearance was significantly delayed. These results align with prior studies demonstrating that disruption of innate immune sensing pathways compromises early neutrophil responses and bacterial clearance. Geddes et al. showed that NOD1-deficient mice infected with *Citrobacter rodentium* displayed impaired induction of IL22, leading to delayed pathogen clearance.^32^ Similarly, Nod1-deficient mice infected with *Clostridium difficile* exhibited increased bacterial translocation, reduced CXCL1 production, impaired neutrophil trafficking, and reduced bacterial clearance, ultimately leading to heightened mortality.^33^ NOD1 KO mice infected with *Salmonella Typhimurium* also displayed defective lamina propria dendritic cell responses, including reduced iNOS/NO production and impaired bacterial clearance.^34^ Together, these studies highlight the crucial role for NOD1 in maintaining the host response to enteric infections. In the current study, reduced clearance of EPEC observed in Nod1^ΔIEC^ mice likely results from impaired chemokine signaling, as evidenced by reduced *Ccl2, Cxcl1,* and *Ccr2* expression in the distal ileum. These findings were further supported by flow cytometry data, whereby monocyte and macrophage frequencies increased in WT EPEC-infected mice, due to their recruitment during early enteric infection, were blunted in Nod1^ΔIEC^ EPEC-infected mice, indicating that loss of epithelial NOD1 impairs immune cell trafficking and may contribute to delayed bacterial clearance. Collectively, these studies support NOD1 as a critical epithelial sensor initiating antimicrobial responses–including cytokine/chemokine secretion, myeloid cell recruitment, and activation of enteric neurons –during neonatal enteric infection. Consistent with this, loss of IEC NOD1 blunts these defenses, delaying pathogen clearance.

Given the delayed bacterial clearance observed in Nod1^ΔIEC^ EPEC-infected mice, mucosal damage was assessed. Histological analysis of the distal ileum revealed significant villus blunting and reduced crypt-villus axis length in WT mice infected with EPEC, but not in Nod1^ΔIEC^ mice. These macroscopic changes likely reflect ongoing epithelial stress and impaired regenerative capacity, as suggested by the significant reductions in Ki67^+^ IEC in these mice. Histological changes after EPEC infection are consistent with prior studies in adult mice pre-treated with streptomycin, where infection induced microvillous effacement, crypt abscesses, neutrophil infiltration, and goblet cell expansion.^13^ To gain insight into whether Nod1^ΔIEC^ mice have reduced injury, impaired initiation of repair, or both, expression of genes critical to these processes were assessed. Neonatal EPEC infection induced a robust innate and adaptive mucosal immune response in the distal ileum of WT mice with increased expression of pro-inflammatory cytokines (*Il1β*, *Il6*, *Il12*) and epithelial regulatory genes (*Il22*, *Tgf*β*1*, *Il10*), which are consistent with our prior study.^8^ The concurrent upregulation of PRR, including *Tlr4* and *Nod2,* supports a model in which select microbial product can activate innate immune pathways in IEC, amplifying local immune activation.

The absence of this immune response in Nod1^ΔIEC^ mice strongly implicates epithelial NOD1 as a necessary upstream regulator of EPEC-induced cytokine and PRR gene expression. The ENS continues to develop and mature during early life, making it susceptible to the effects of neonatal enteric infection.^35^ Here, we found that neonatal EPEC infection caused significant neuronal loss in the myenteric plexus of the distal ileum at P14. This is consistent with enteric pathogens inducing inflammation-driven neurotoxicity in the gut, where cytokines such as *Tnfα* and *Il1β* promote oxidative stress, glial activation, and loss of myenteric neurons, particularly in the developing ENS.^36^ Moreover, neuronal loss was biased toward excitatory neurons and persisted for months post-infection, resulting in sustained GI dysmotility following *Salmonella* infection.^36^ Neuronal loss coupled with glial activation, as demonstrated by increased S100b expression, during neonatal infection may permanently reprogram neural circuits and impair immune–neural crosstalk, providing a mechanistic basis for the long-term GI dysfunction and altered barrier integrity observed in adult WT mice. Together, these data identify the ENS as a critical and vulnerable target of early-life enteric infection and highlight the need for therapeutic strategies to protect or restore enteric neurons during neonatal inflammation.

GI inflammation seen in neonatal WT mice following EPEC infection persisted into adulthood, well past clearance of the pathogen, and was coupled with GI pathophysiology.^8^ Ussing chamber studies indicated increased ileal intestinal permeability by FITC-dextran flux and transepithelial conductance in adult WT, but not Nod1^ΔIEC^ mice previously infected with EPEC. These data align with findings of ongoing elevated inflammatory gene expression (*Il1β*, *Il6*, *Il12*) and barrier repair markers (*Il22*, *Slc26a3*) in adult WT mice neonatally infected with EPEC due to persistent inflammation and compensatory repair mechanisms. Activation of NOD1 may exacerbate infection-induced damage by amplifying NF-κB–mediated inflammatory signaling and stress responses in IEC, leading to persistent epithelial dysfunction. These findings suggest that during infection, IEC NOD1 activation can cause chronic mucosal barrier leakiness and impaired tight junction dynamics, which are known contributors to ongoing mucosal inflammation and systemic immune activation.^37^ These results suggest a lasting impact of neonatal enteric infection on intestinal physiology, and they reinforce NOD1 as a candidate target for interventions aimed at preserving epithelial function following neonatal infectious insult.

Behavioral studies previously revealed that neonatal EPEC infection has long-lasting consequences on cognitive function, specifically impairments in recognition memory in adulthood as demonstrated by the NOR task.^8^ This reduction in recognition memory was confirmed in WT mice in our current study and was significantly attenuated in Nod1^ΔIEC^ mice. These data strongly implicate epithelial NOD1 in causing the impaired cognitive sequelae following early-life infection. Normal anxiety-like behavior and locomotor activity in WT and Nod1^ΔIEC^ mice following neonatal infection suggests a unique mechanism driving deficits in hippocampal-dependent cognition, as seen previously.^8^ Cognitive deficits previously found in adult mice following neonatal EPEC infection were coupled with hippocampal neuroinflammation, highlighting the impacts of enteric bacterial infection on the central nervous system.(Henessey) Here we similarly demonstrate increased hippocampal *Iba1* expression, a marker of microglia, coupled with increased cytokines (*Il6*, *Il1β*, *Tnfα, Il10)* and PRR *(Tlr4)* in WT mice. As seen in the GI tract, no evidence of neuroinflammation was present in Nod1^ΔIEC^ mice, implicating epithelial NOD1 in modulating downstream CNS impairments following neonatal enteric infection. These findings mimic those seen in a viral encephalitis model, NOD1 deficiency dampened microglial activation and pro-inflammatory cytokine responses, suggesting that NOD1 can directly influence neuroinflammatory tone in the brain following infection.^38^ However, the absence of hippocampal responses in Nod1^ΔIEC^ mice suggests that a reduced gut-derived inflammatory signaling, rather than a loss of central NOD1 function, is mediating these effects. Prior CNS studies demonstrate that microglial NOD1 can amplify neuroinflammation when appropriately stimulated^38,39^, potentially allowing for IEC NOD1-driven signals to converge on central NOD1-dependent pathways. Behavioral deficits and neuroinflammation were also coupled with evidence of increased hippocampal neurogenesis. Increased hippocampal *Bdnf* expression and increased Ki67⁺ proliferative cells and DCX⁺ immature neurons were found in WT, but not Nod1^ΔIEC^ mice, neonatally infected with EPEC. Together, these findings support a model in which neonatal enteric infection causes hippocampal neuroinflammation and exerts long-lasting consequences on neurogenesis and disrupted hippocampal plasticity, detrimentally impacting cognitive function in a Nod1-dependent manner. Collectively, these findings underscore the potential for early microbial insults to exert long-lasting effects on neuroimmune homeostasis and highlight epithelial NOD1 as a mechanistic bridge between GI infection and brain inflammation.

NLRs are important in regulating serotonin signaling in both the gut and the brain in response to stress^19^ likely via both ENS and CNS neurons. Here we found persistent alterations in serotonin signaling in the distal ileum, including increased *Tph2*, *5Htr2c*, and *5Htr4* expression at both P14 and in adult WT EPEC-infected mice, but not Nod1^ΔIEC^ mice. Similar impairments in 5-HT signaling were found in the adult WT hippocampus, including increased expression of *Tph2, Sert,* and *5Htr2c*, whereas *5Htr4* was unaffected. Together, these findings suggest that serotonergic dysregulation following neonatal enteric infection occurs both locally in the gut and distally within the CNS, and support the idea that serotonergic dysregulation may be caused by chronic inflammation or MGB axis perturbation in neonatal life.^40,41^ Serotonin is increasingly recognized as a modulator of immune tone, glial reactivity, and neurogenesis^41,42^; its dysregulation likely contributes to the cognitive and plasticity deficits we observed. Altered HPA-axis activation was also identified, including increased *Crfr2* expression in the distal ileum (neonatal and adult) and hippocampus (adult), which was coupled with increased serum corticosterone levels in WT mice. Importantly, these serotonergic and HPA-axis alterations were absent in Nod1^ΔIEC^ mice, indicating that epithelial NOD1 is required to propagate signals following neonatal infection to systemic neuroendocrine pathways. Together, these findings strongly support the idea that neonatal EPEC infection reprograms host serotonergic signaling via IEC NOD1, ultimately impairing stress axis regulation and hippocampal function. IEC NOD1 therefore represents a key regulator of long-term serotonergic tone across the gut and brain, linking early-life EPEC infection to sustained alterations in neuroendocrine and behavioral outcomes.

Probiotics can beneficially regulate the MGB axis, particularly following infection with an enteric pathogen.^43^ Despite these benefits, it remains unclear by which mechanism these effects are mediated.^44^ In contrast to NOD1, which provides innate immunity and inflammatory programming following infection, activation of NOD2 has been found to be protective against intestinal inflammation.^45^ Using LC/MS, we identified a heterogeneous composition of muropeptides isolated from probiotic Lactobacillus sp, reflecting the physiological diversity of muropeptides generated during bacterial cell wall turnover.

Importantly, we identified the presence of the immunomodulatory M-Tri-Lys peptide, a NOD2 ligand, previously reported to ameliorate chemically induced colitis in mice.^46^ We found that probiotic *Lactobacillus* sp. derived muropeptides did not elicit pro-inflammatory responses, but rather induced a robust *Il10* response in cultured splenocytes. Supplementation of neonatal mice with probiotic-derived muropeptides prevented EPEC-induced ileal pro-inflammatory cytokine (*Il6*, *Tnfα*, *Il1β*) expression while preserving anti-inflammatory (*Il10*) expression at P14. This is in keeping with prior evidence that PGN/muropeptides reduced colitis severity by NOD2-dependent IL-10 production, and T regulatory development by dendritic cells.^46^ Administration of probiotic muropeptides also reduced expression of PRR, and chemokine ligand/receptor expression, further suggesting reduced pro-inflammatory microbial sensing pathways and immune cell recruitment following enteric infection.

Interestingly, probiotic muropeptide supplementation did not restore levels of *Muc2* expression in EPEC-infected mice, nor decrease the elevated numbers of goblet cells to LB control levels. This may serve a protective role following infection^47^ and importantly it also indicates that muropeptide supplementation did not simply block all host responses to enteric infection but rather supported a beneficial response. These data suggest that probiotic muropeptide administration could potentially improve mucus layer functionality—an effect that reinforces the mucosal barrier against luminal pathogens. Lastly, the ability of muropeptides to protect enteric nerves from death following enteric infection is highly exciting and novel, proposing a key mechanism by which host-microbe interactions may ensure maintenance of GI physiology, and limit symptoms including diarrhea. Together, our data identifies probiotic muropeptides as beneficial host-directed immunomodulators of enteric infection and by extension infection-induced gut-brain deficits. By tuning epithelial PRR signaling, restraining immune cell trafficking, reinforcing barrier integrity, and protecting enteric neurons, muropeptides offer a targeted approach to limit inflammation in the immature gut—one with translational potential for preventing long-term neuroimmune sequelae of early-life enteric infections.

In conclusion, neonatal EPEC infection causes MGB axis deficits through neuroimmune deficits mediated via IEC NOD1, which can be beneficially modulated by probiotic muropeptides. Together, these studies highlight the important role of NLRs in coordinating the host neuroimmune response to infection and subsequently driving gut-brain deficits. Excitingly, this study identifies a potentially novel therapeutic strategy for pediatric enteric bacterial infections.

## Abbreviations

5-HT: Serotonin
AMPs: Antimicrobial peptides
CCh: Carbachol
CNS: Central nervous system
DC: Dendritic cell
ENS: Enteric nervous system
GABA: γ-Aminobutyric acid
GI: Gastrointestinal
IBD: Inflammatory bowel disease
IEC: Intestinal epithelial cell
IFNγ: Interferon-gamma
IL: Interleukin
KO: Knockout
DKO: Double knockout
LPS: Lipopolysaccharide
MGB: Microbiota-gut-brain
MUC2: Mucin 2
NF-κB: Nuclear factor kappa-B
NLR: NOD-like receptor
NOD: Nucleotide-binding oligomerization domain protein
PAMP: Pathogen-associated molecular pattern
PGN: Peptidoglycan
PRR: Pattern recognition receptor
T3SS: Type 3 secretion system
TLR: Toll-like receptor
TNFα: Tumor necrosis factor-alpha
Treg: Regulatory T cells

## ACKNOWLEDGEMENTS

This work was funded in part by R21MH108154 (MGG), R01AT009365 (MGG), T32MH073124 (JRG), and T32AI060555 (MC). The authors wish to acknowledge the support of the UC Davis Comprehensive Cancer Center Flow Cytometry Shared Resource, supported by the NIH (P30CA093373).

OO was an officer of the United States Space Force during her graduate training at UC Davis. The views expressed in this article are those of the author and do not reflect the official policy or position of the United States Air Force, the Department of Defense, or the U.S. Government

The authors wish to thank the UC Davis Health District Advanced Imaging Facility (HDAIF) for help with immunofluorescence. ANNA-1 antibody for neuronal staining was kindly gifted by Dr. Vanda Lennon (Mayo Clinic).

